# A metabolically resistant spexin analogue, LIT-01-144, induces potent non-opioid peripheral antinociception in persistent pain via activation of GALR2

**DOI:** 10.64898/2026.02.19.706558

**Authors:** Yann Berthomé, Glenn-Marie Le Coz, Valérie Utard, Qiuxiang Gu, Rosine Fellmann-Clauss, Nathalie Petit-Demoulière, Raphaëlle Quillet, Claire Gavériaux-Ruff, Sridévi Ramanoudjame, Lucie Estéoulle, Nicolas Humbert, François Daubeuf, Patrick Gizzi, Stéphanie Riché, Xavier Leroy, Dominique Bonnet, Frédéric Simonin

## Abstract

Chronic pain affects a significant portion of the global population and imposes substantial clinical and socioeconomic burdens. Current treatments mainly rely on opioid analgesics, which carry serious risks of dependence and misuse, underscoring the urgent need for alternative therapeutic strategies. Galanin receptors (GALR1-3) are known to be involved in modulating pain, yet their specific roles remain poorly understood due to the lack of receptor subtype-selective ligands. Recently, spexin has been identified as an endogenous peptide that selectively activates GALR2 and GALR3, offering a new scaffold for developing pharmacological tools targeting these receptor subtypes. In this study, we report the design and characterization of a modified spexin analog, LIT-01-144, engineered through N-terminal functionalization with a fluorocarbon chain to improve metabolic stability while preserving receptor selectivity. In vitro assays showed that LIT-01-144 has high potency at GALR2 and GALR3, with minimal activity at GALR1. Pharmacokinetic studies revealed a significantly longer plasma half-life compared to native spexin and no central nervous system penetration. In mice, intracerebroventricular administration of LIT-01-144 produced strong antinociceptive effects at doses ten times lower than spexin. While systemic administration showed no notable antinociception in naïve animals, LIT-01-144 significantly reduced pain responses in a mouse model of persistent inflammatory pain induced by complete Freund’s adjuvant (CFA). This antinociceptive activity was specifically mediated through GALR2 and was independent of opioid receptor pathways. In situ hybridization further showed an increase in Galr2-positive neurons in dorsal root ganglia of inflamed mice. Overall, these findings highlight GALR2 as a promising peripheral target for developing non-opioid analgesics and demonstrate the potential of LIT-01-144 as a valuable tool for understanding GALR2-mediated mechanisms of pain modulation.

## Introduction

Chronic pain is one of the most significant conditions in terms of clinical and socioeconomic impact. Approximately 20% of U.S. adults experience chronic pain (Dahlhamer et al., 2018). Similarly, in Europe, 25%–35% of adults are affected (Breivik et al., 2013). Opiates remain the gold standard for pain management despite numerous adverse side effects, including constipation, nausea, sedation, respiratory depression, tolerance and dependence. The latter three are particularly concerning, as they contribute to opioid misuse, self-prescription, overdose, and death (Busserolles et al., 2020; Mattson et al., 2021), highlighting the urgent need for new therapeutic strategies.

The galanin receptor (GALR) family is a well-characterized subgroup of three G protein-coupled receptors (GPCRs) that bind the endogenous peptide galanin and are involved in the modulation of different functions including feeding, energy homeostasis, osmotic regulation, water intake and pain (Lang et al., 2015). Despite extensive research, the precise role of each receptor in the modulation of pain remains unclear, largely due to the lack of selective agonists or antagonists (Hulse et al., 2011; Fonseca-Rodrigues et al., 2022). GALR1, the most abundant receptor, is widely distributed in the central nervous system (CNS) and peripheral nervous system including the spinal dorsal horn and dorsal root ganglia (DRG), and has been shown to display antinociceptive properties (Lang et al., 2015). Indeed, selective destruction of GALR1-containing dorsal horn neurons leads to heat hypoalgesia (Lemons and Wiley, 2011) and GALR1-knockout (KO) mice exhibit increased thermal and mechanical hypersensitivity, particularly following tissue injury or nerve damage (Blakeman et al., 2003; Malkmus et al., 2005). Although GALR2 is predominantly expressed in the brain and DRG (O’Donnell et al., 1999), its contribution to nociceptive modulation remains complex and contentious, as both pro-nociceptive and antinociceptive effects have been documented. For example, intra-plantar administration of the GALR2 agonist M1896 in a rat neuropathic pain model exacerbated tactile allodynia, while the GALR2 antagonist M871 reversed this effect (Chen et al., 2018). Conversely, administration of NAX 409-9, a PEG-Galanin GALR2-preferring agonist, produced an antinociceptive response in models of inflammatory and neuropathic pain in rats (Metcalf et al., 2015). GALR2-KO mice exhibit reduced neuropathic and inflammatory pain responses, however interpretation of these findings is complicated by the developmental deficits of DRG neurons in these animals (Hobson et al., 2006). GALR3 has been less studied due to its lower expression. Its distribution in the CNS is sparse and highly restricted (Mennicken et al., 2002). While its precise role remains unclear, one study has shown increased paw inflammation in GALR3 KO mice in a murine model of autoimmune arthritis but mechanical allodynia was similar to that of wild-type mice (Botz et al., 2016).

Recently, a new endogenous neuropeptide named spexin (alias neuropeptide Q) was discovered by bioinformatic analysis (Mirabeau et al., 2007; Sonmez et al., 2009) and was further shown to selectively activate GALR2/R3 but not GALR1 (Kim et al., 2014). Intracerebroventricular (icv) administration of spexin in naïve mice induced thermal antinociception, an effect that was not blocked by the opioid antagonist naltrexone (Toll et al., 2012). Additionally, icv injection of spexin in ovariectomized rats reduced pain sensitivity in the formalin test, indicating a potential role in the modulation of inflammatory pain (Moazen et al., 2018). Similarly, spexin icv administration in mice displayed antinociceptive activity in the formalin test and in the acetic acid-induced writhing test (Lv et al., 2019a). However, the precise role of spexin in peripheral pain modulation and its interaction with GALR2/R3 remains unclear (Lv et al., 2019b).

In this study, we describe the design, synthesis and characterization of a novel pharmacological tool derived from spexin, LIT-01-144, to study the role of GALR2/R3 *in vivo*. This compound exhibits improved metabolic stability and agonist activity toward GALR2/R3 as compared to spexin while retaining a similarly very low affinity toward GALR1. In mice with persistent inflammatory pain, LIT-01-144 displayed antinociceptive activity following peripheral administration at doses 10 to 100-fold lower than spexin. This activity was associated with an increase in the number of neurons that expressed *Galr2* in DRG and was independent of GALR3 and opioid receptors. These results provide evidence for a major role of spexin and GALR2 in the modulation of nociception in an opioid-insensitive manner and further highlight the therapeutic potential of GALR2-targeting compounds in pain management.

## Material and methods

### Compounds

Human galanin (hGal) and spexin peptides (> 95% purity) were purchased from Genecust®. SNAP-37899 was obtained from MedChemExpress and naltrexone from Sigma-Aldrich. LIT-01-144 and LIT-01-128 were synthesized by Fmoc/tBu solid phase peptide synthesis from commercially available Rink amide resin, Fmoc-amino acids and fluorocarbon chains. Crude peptides were purified by semi-preparative reversed-phase high-performance liquid chromatography (RP-HPLC) on a Gilson PLC2020 system. The identity and purity of the peptides were assessed by analytical RP-HPLC and by LC-MS. All the tested compounds displayed purity > 95%. More details are provided in the supplementary methods.

### Plasma stability assay

Plasmatic stability of spexin and LIT-01-144 (0.25 μM) was determined in CD1 mouse plasma. Plasma was prewarmed at 37 °C for 5 min before the addition of the tested compound. The mixture was incubated at 37 °C for 24 h. An aliquot of the incubation mixture was transferred to ice-cold MeCN (3 volumes) at defined times (15, 30, 60, 180, 360 and 1440 min). Samples were frozen at -80 °C, unfrozen, mixed, and centrifuged. Supernatants were collected and analyzed by UPLC-MS/MS without further dilution. The percentage of remaining peptide at each time point was determined by monitoring the peak area of the chromatogram. Half-life (*t*_1/2_) was estimated from the slope of the initial linear range of the logarithmic curve of the remaining compound (%), assuming that the degradation follows a first-order kinetic.

### Compounds solubilization for in vitro assays

Stock solutions (10 mM) of galanin, spexin and fluorospexin derivatives were prepared in DMSO molecular biology grade (Sigma). For fluorospexin derivatives solutions were subjected to 3 cycles of 30s sonication and 30s vortex, then diluted in HEPES/1% BSA (10 mM HEPES, 0.4 mM NaH_2_PO_4_, 137.5 mM NaCl, 1.25 mM MgCl_2_, 1.25 mM CaCl_2_, 6 mM KCl, 5.6 mM glucose, 1% (W/V) BSA, pH = 7.4) physiological buffer.

### Glosensor cAMP assay

cAMP accumulation assay was essentially done as described (Roeckel et al., 2017) with the following modifications: HEK293 cells stably expressing GalR1 cultivated at 37 °C and 5% of CO_2_ in DMEM (4.5 g/l glucose, Gibco), supplemented with 10% of fetal calf serum (PAN Biotech), 100 UI/ml of penicillin and 100 μg/ml of streptomycin (Gibco) were used, with additional transient expression of pGloSensor-22F using lipofectamine 2000 (Invitrogen) as transfection reagent. The cAMP responses were measured in the presence of D-luciferin (1 mM; Synchem). All peptides and test compounds were incubated in the presence of IBMX (0.1 mM; Merck) for 10 min before inducing cAMP production by forskolin (5 µM; Merck). The maximum luminescence levels were plotted depending on the agonist concentration using Graphpad/Prism software allowing the determination of the EC_50_ value of the agonist.

### Calcium mobilization assay

HEK293 cells stably expressing the human GALR2 were cultivated at 37 °C and 5% of CO_2_ in DMEM (4.5 g/l glucose, Gibco), supplemented with 10% of fetal calf serum (PAN Biotech), 100 UI/ml of penicillin and 100 μg/ml of streptomycin (Gibco). For GALR3, HEK293 cells stably expressing GqiTop (Selvam et al., 2010) cultivated at 37 °C and 5% of CO_2_ in MEM (Gibco), supplemented with 10% of fetal calf serum (PAN Biotech), 100 UI/ml of penicillin and 100 μg/ml of streptomycin (Gibco) were transfected using lipofectamine 2000 to afford transient expression of GALR3. The day of the experiment, for GALR2, cells were incubated with 2 µM Fluo-4 AM (Molecular Probes) for 1 h at 37 °C, rinsed, and distributed in a black 96-well plate (about 500 000 cells/well in HEPES/1% BSA). For GALR3, cells were incubated with 2 µM Calbryte 520 AM (AAT Bioquest) for 1 h at 37 °C and 15 min at room temperature, rinsed, and distributed in a black 96-well plate (about 500 000 cells/well in HEPES/1% BSA). Agonist-evoked increase in intracellular calcium was recorded over time (5 sec intervals over 220 sec) at 37 °C through fluorescence emission at 520 nm (excitation at 494 nm) in a FlexStation 3 plate reader (Molecular Device). Peak response amplitudes were normalized to basal and maximal (cells permeabilized with 150 μM digitonin) fluorescence levels. Finally, the amplitudes obtained after this normalization were plotted depending on the agonist concentration using Graphpad/Prism software allowing the determination of the EC_50_ value of the agonist.

### Animals

Animal experiments were performed on adult C57BL/6N (25-30 g; Janvier Labs, France) or CD1 male mice (35 g; Janvier Labs, France). Animals were housed in groups of three to five per cage and kept under a 12 h/12 h light/dark cycle at 21 ± 1 °C with *ad libitum* access to food and water. Experiments were performed during the light-on phase of the cycle. Mice were habituated to the testing room and equipment and handled for two weeks before starting behavioral experiments. All experiments were carried out in strict accordance with the European guidelines for the care of laboratory animals (European Communities Council Directive 2010/63/EU), approved by the local ethical committee and authorized by the French Ministry for Research (authorization numbers: #14586, #19987). All efforts were made to minimize animal discomfort and to reduce the number of animals used.

### In situ hybridization of dorsal root ganglia

Dorsal root ganglia of naïve and CFA-treated mice were collected and directly incorporated in optimum cutting temperature medium (Sakura Fintek) then frozen in dry ice. Tissues were cryo-sectioned (14 µm; -20 °C) and collected on Superfrost Plus glass slides (Fisher). Commercially available RNAscope Multiplex fluorescent v2 kits and probes were purchased directly from Advanced Cell Diagnostics. In situ hybridization probes were designed against mouse GALR2 (bp 463-1539; NM_010254.4; C1) and mouse NeuN (Mm-Rbfox3 313311-C2). Fluorophore was assigned to the C1 or C2 channel depending on the selected tyramide signal amplification (TSA) plus fluorophore (Perkin Elmer) diluted at the optimized concentration (GALR2-C1: TSA Plus Cyanine 3 1/1500, C2: TSA Plus Cyanine 5 1/1500). In situ hybridization was performed according to the manufacturer’s recommendation with slight optimization. Briefly, a post-fixation step (freshly prepared prechilled 4% paraformaldehyde at 4 °C during 1h) was followed by 3 washes with PBS (3 x 1 min at room temperature) and dehydration with ethanol bath (50%, 70%, 100%, 100%, 5 min each, room temperature). After slides pretreatment with Protease Pretreat 4 (Advanced Cell Diagnostics) at room temperature for 20 min, hybridization was performed at 40 °C for 2 h in a HybEZ oven (Advanced Cell Diagnostics). Following wash and amplification steps, DAPI (4’,6-diamidino-2-phenylindole; Molecular Probes) was used to label nuclei and Prolong Gold antifade reagent (Molecular Probes) was used to mount coverslips. Whole images were acquired using Hamamatsu NanoZoomer S60 digital slide scanner (40x) and analyzed with the NDPview software. Data presented were obtained from at least 6 DRGs obtained from 3 mice for each group.

### Pharmacokinetics of spexin and LIT-01-144

6 weeks old CD1 mice were injected with 10 mg/kg (ip) of spexin (solubilized in saline) or LIT-01-144 (solubilized in H_2_O + 5% DMSO + 1% tween 80). For animals injected with spexin (n = 3), blood samples (30 μl) were collected by tail incision at different time points (2, 4, 7, 9, 12 and 15 min). Animals injected with LIT-01-144 (n = 3 for each time point) were anesthetized by intraperitoneal administration of a mixture of ketamine 100 mg/kg (Imalgène 1000) and xylazine 10 mg/kg (Rompun), and blood samples (400 μl) were collected by an intracardiac puncture at different time points (3, 15, 30, 60, 180 and 360 min). All the samples were placed into EDTA-coated tubes, centrifuged at 4 °C, 10 000 g for 10 min and the plasma was stored at -80 °C. Plasma samples were analyzed by UPLC-MS/MS (LC-MS 9030. Shimadzu, electrospray ion source) using a C18 column (Phenomenex 2.6 μm Kinetex, 50 X 2.1 mm) and a linear gradient (5% to 95% in 1.2 min, flow rate of 0.5 ml/min) of MeCN with 0.05% formic acid (v/v) in H_2_O with 0.05% formic acid (v/v). Plasma samples were processed before analysis as follows: 13 μl (spexin) or 400 µl (LIT-01-144) of plasma samples were mixed with 33 µl or 1ml (respectively) of MeCN for protein precipitation and compound extraction. Samples were vortex-stirred for 5 min, sonicated for 1 min, and centrifuged at 15,000 g for 5 min at 16 °C. Peptides were quantified in the supernatant by integrating the area under the peaks, and normalization was based on the standard and compared to standard curves. Standard curves were obtained by analyzing known peptide quantities that were dissolved in plasma and processed and analyzed using the same procedure. Pharmacokinetics parameters were calculated using a one-compartment model.

### Brain concentration determination

Spexin and LIT-01-144 were injected at a dose of 10 mg/kg (ip) in C57BL/6N male mice. Brain samples were collected at different time points (see Table S1). Before recovery of the samples, mice were anesthetized with a solution of ketamine (100 mg/kg, Imalgène 1000) / xylazine (10 mg/kg, Rompun) at 37 °C, injected intraperitoneally (10 ml/kg), followed by a subcutaneous (sc) injection of lidocaine 10 mg/kg at the incision site 5 minutes beforehand. The chest was opened and mice were sacrificed by intracardiac puncture and brains were collected after perfusion of 50 ml cold PBS with a peristaltic pump (10 ml/min). Brains were homogenized in a volume of water corresponding to its mass using an Ultra-Turrax, frozen, and stored at -80 °C. For the analyses, two volumes of acetonitrile were added for each homogenized brain. Samples were vortexed for 3 minutes, and placed in an ultrasonic bath for 3 minutes. Precipitated proteins and solid residues were centrifuged (15 000 g, 5 min at 16 °C) and the supernatants were transferred to a microplate for LC-MS/MS analysis. All samples were analyzed using a UPLC coupled to a Shimadzu LC-MS 8030 triple quadrupole. Standards for calibration were prepared with brains from untreated mice. Samples were analyzed twice in the same series of injections.

### Assessment of thermal nociception

The thermal nociceptive threshold of the animals was assessed using the tail immersion test as previously described (Elhabazi et al., 2014). Briefly, mice were restrained in a grid pocket and two-thirds of their tails were immersed in a thermostatic water-bath heated at 47.5 ± 0.5 °C. The latency (in sec) for tail withdrawal from hot water was taken as a measure of the nociceptive response. In the absence of any nociceptive reaction, a cut-off value of 25 sec was set to prevent tissue damage.

### Assessment of mechanical nociception

The mechanical nociceptive threshold of the animals was assessed using the tail pressure test. Mice were restrained in a grid pocket, with the tail sticking out. Using an instrumented algometer (BIO-RP-WRS, Bioseb), increasing pressure was exerted on the first third of the tail on the proximal side, until the first perceptible reaction from the mouse (tail withdrawal, body movement, vocalization, etc.). The value, in grams, was noted as a measure of the nociceptive response. In the absence of any nociceptive reaction, a cut-off value of 3 x the basal threshold was set to prevent tissue damage.

### Central and peripheral antinociceptive activity of spexin and LIT-01-144 in naïve mice

Intracerebroventricular injections (5 µl) were performed under isoflurane anesthesia with a Hamilton syringe into C57BL/6N male mice. Nociceptive latencies were determined before (baseline) and after drug or saline (control) administration (i.e. 15, 30, 45, 60, 90 and 120 min), n = 6-8. For the evaluation of peripheral activity, compounds were injected intraperitoneally in C57BL/6J male mice and nociceptive latencies were determined before (baseline) and after drug or saline (control) administration (i.e. 15, 30, 60, 90, 120, 180, 240, 300, 360, and 1440 min), n = 7 per group.

### Inflammatory pain model

Inflammatory pain in C57BL/6N male mice was induced by injecting subcutaneously at 1 cm from the tip of the tail 20 μl of Complete Freund’s Adjuvant (CFA, Merck) and the development of hyperalgesia was followed with the tail immersion test. Compounds were injected (ip) when hyperalgesia was maximal at day 2 or 3 depending on the experiment and thermal nociceptive threshold of the animals was determined before (baseline) and after drug or saline (control) administrations during several hours/days until they return to their basal nociceptive value (n = 6-8).

### Data analysis

Data were analyzed and graphs were created using GraphPad Prism software. Antinociception was quantified as the area under the curve (AUC) calculated by the trapezoidal method. Data were analyzed using one-way or two-way analysis of variance (ANOVA). Post-hoc analyses were performed with Bonferroni tests. The level of significance was set at p < 0.05.

## Results

### Design, synthesis and functional evaluation of new spexin conjugates

In this study, we sought to develop metabolically stable spexin conjugates that retain the same pharmacological profile than endogenous spexin toward GALRs in order to investigate the role of GALR2/R3 in the modulation of nociception. In previous works, we developed an original strategy that consist in the modification of bioactive peptides with a fluorocarbon chain (FC), which we successfully applied to apelin peptides to obtain metabolically stable candidates for the treatment of hyponatremia, cardiovascular diseases or multiple sclerosis (Gerbier et al., 2017; Flahault et al., 2021; Birmpili et al., 2024; Girault-Sotias et al., 2024). We thus decided to extend this strategy to spexin by incorporating a FC at the C-terminus or N-terminus part of this peptide, resulting in the corresponding fluorinated spexin analogs (LIT-01-128 and LIT-01-144). These peptides were obtained following a solid-phase Fmoc/tBu strategy (supplementary methods) and their sequences are reported in **Table 1**.

**Table 1.** Structures of fluorinated peptides developed to improve spexin *in vivo* properties.

| Peptides | N-ter | 1 | 2 | 3 | 4 | 5 | 6 | 7 | 8 | 9 | 10 | 11 | 12 | 13 | 14 | 15 | C-ter |
| --- | --- | --- | --- | --- | --- | --- | --- | --- | --- | --- | --- | --- | --- | --- | --- | --- | --- |
| Spexin | H | N | W | T | P | Q | A | M | L | Y | L | K | G | A | Q | - | NH<sub>2</sub> |
| LIT-01-128 | H | N | W | T | P | Q | A | M | L | Y | L | K | G | A | Q | K(FC*) | NH<sub>2</sub> |
| LIT-01-144 | FC* | N | W | T | P | Q | A | M | L | Y | L | K | G | A | Q | - | NH<sub>2</sub> |

\* Fluorocarbon Chain (FC) = CF<sub>3</sub>(CF<sub>2</sub>)<sub>7</sub>(CH<sub>2</sub>)<sub>2</sub>C(O)-

To determine the impact of FC incorporation and location on the pharmacological properties of these spexin derivatives, we first evaluated the *in vitro* activity of LIT-01-128 and LIT-01-144 in comparison to spexin and galanin on HEK293 cells expressing GALR1, GALR2 and GALR3 (**Fig. 1 and Table 2**). As previously described (Kim et al., 2014) spexin displayed almost no agonist activity on GALR1 (EC_50_ > 10 000 nM) as compared to galanin that showed sub-nanomolar potency (EC_50_ = 0.6 ± 0.1 nM). Similarly to spexin, LIT-01-128 and LIT-01-144 displayed poor activity at GALR1 (EC_50_ > 10 000 nM) while they displayed full agonist activity at both GALR2 and GALR3 (**Fig. 1**, **Table 2**). LIT-01-144 was the most potent at both GALR2 and GALR3 with EC_50_ values of (4.2 ± 0.9 nM and 3.2 ± 0.7 nM, respectively) that were 10-fold lower than endogenous spexin (53 ± 5 nM and 37 ± 9 nM, respectively). These results indicated that the N-terminal position of spexin was most suited for the introduction of a FC than the C-terminal position while maintaining the same selectivity profile as spexin. Based on these data we therefore decided to characterize further LIT-01-144.

**Figure 1.**
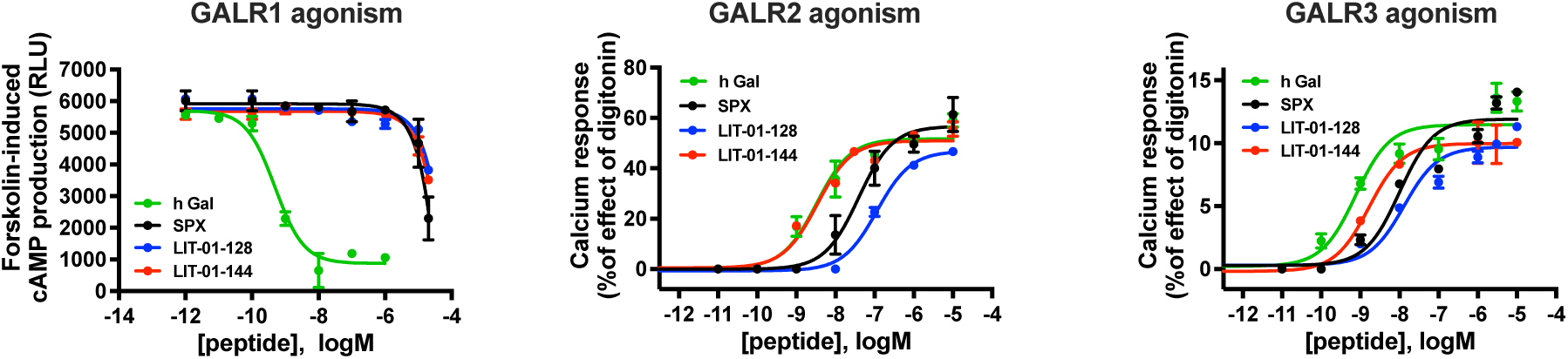
In vitro functional characterization of human galanin, spexin, and fluorospexin ligands at human GALR1, GALR2, and GALR3. The activity of the different compounds was evaluated on HEK293-Glo22F-hGALR1 cells in inhibition of cAMP production assay, on HEK293-hGALR2 and HEK293-GqiTOP-GALR3 cells in calcium mobilization assay. The data shown are representative experiments of at least three independent experiments performed in duplicate.

**Table 2.** Half-maximal effective concentration (EC_50_) values of human galanin, spexin and fluorospexin derivatives for human GALR1, GALR2 and GALR3.

| | $EC_{50}$ (nM) | | |
| --- | --- | --- | --- |
|  | GALR1 | GALR2 | GALR3 |
| hGal | $0.6 \pm 0.1$ | $4.4 \pm 0.5$ | $3.9 \pm 0.7$ |
| Spexin | $> 10\ 000$ | $53 \pm 5$ | $37 \pm 9$ |
| LIT-01-128 | $> 10\ 000$ | $115 \pm 27$ | $654 \pm 93$ |
| LIT-01-144 | $> 10\ 000$ | $4.2 \pm 0.9$ | $3.2 \pm 0.7$ |

Data are mean ± SEM of at least three independent experiments performed in duplicate.

### Pharmacokinetic evaluations of spexin and LIT-01-144

We next evaluated the stability of LIT-01-144 in mouse plasma as compared to that of endogenous spexin. The half-life of LIT-01-144 was found to be 432 ± 6 min, corresponding to a 7-fold increase compared to spexin (t_1/2_ = 65 ± 12 min). We then evaluated the pharmacokinetic properties of LIT-01-144 and spexin after ip administration in mice at a dose of 10 mg/kg (**Fig. 2A**). Consistently with our *in vitro* data, LIT-01-144 displayed a half-life of 101 ± 7 min that was 20 times higher than spexin (5.1 ± 1.7 min, **Table 3)** resulting in an exposure that was around 45 times higher than that of spexin (**Fig. 2B**, **Table 3**). Noteworthy, the absorption of LIT-01-144 and spexin was similar with C_max_ values of 0.20 ± 0.02 nmol/mL and 0.13 ± 0.05 nmol/mL that were reached 30 min and 3 min following their administration, respectively (**Table 3**). These results confirmed that LIT-01-144 was suitable for further *in vivo* evaluation. In addition, we evaluated the susceptibility of spexin and LIT-01-144 to cross the blood brain barrier after ip injection in mice at a dose of 10 mg/kg. Our data showed no detectable traces of spexin and LIT-01-144 in the brain of animals 5, 30 and 360 min after their administration, indicating that these compounds could not cross the blood brain barrier (**Table S1**).

**Figure 2.**
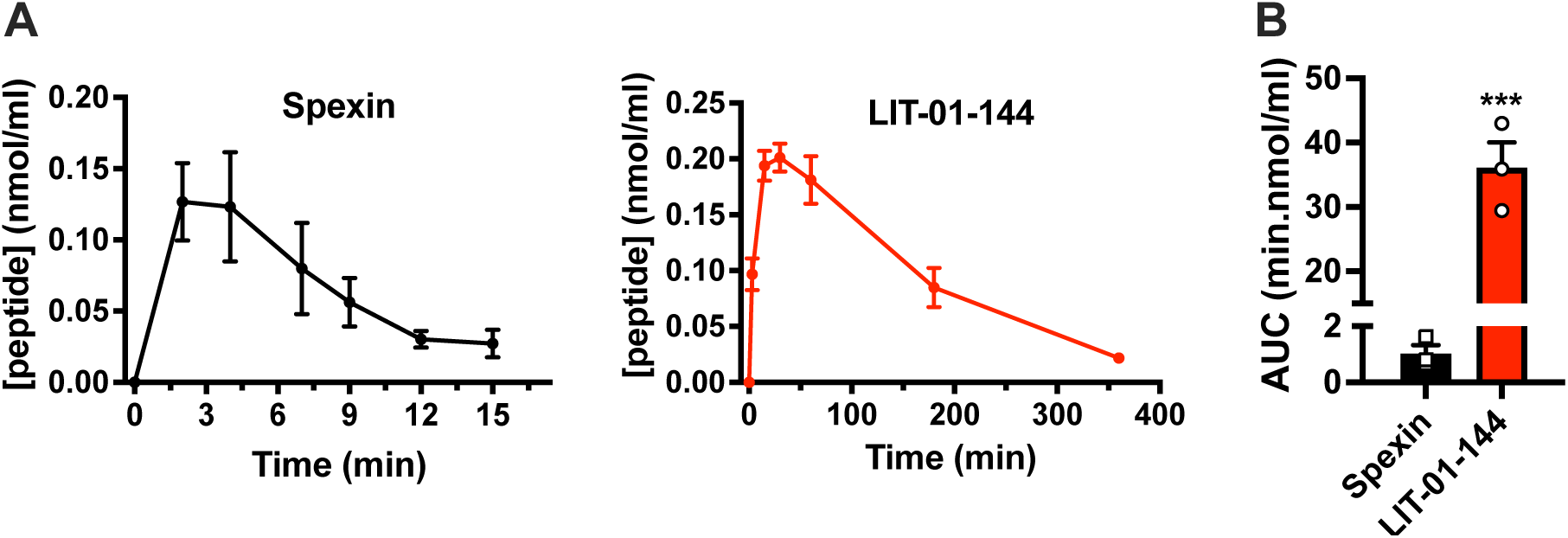
Pharmacokinetics of spexin and LIT-01-144 after ip administration in mice. A) Spexin and LIT-01-144 were administered at 10 mg/kg (ip) and plasma concentrations were determined using mass spectrometry. B) Comparison of the exposition between spexin and LIT-01-144 as area-under-the-curve (AUC) values over 15 min for spexin and 360 min for LIT-01-144. Data are presented as mean ± SEM (n = 3 for spexin and n = 3 per time point for LIT-01-144). Groups were compared using unpaired t-test ***p<0.001.

**Table 3.**
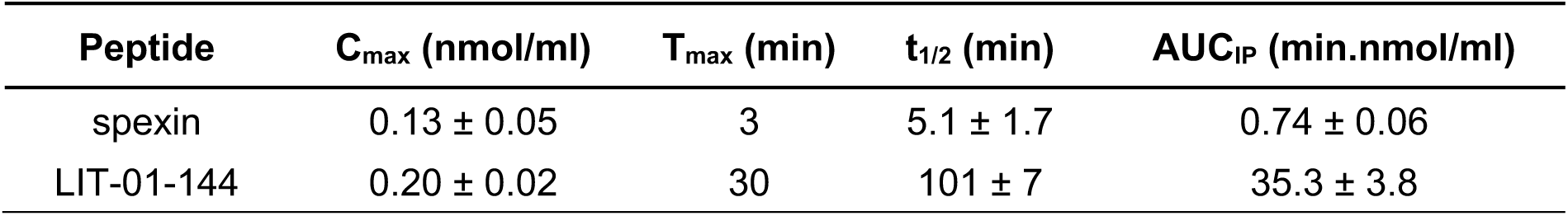
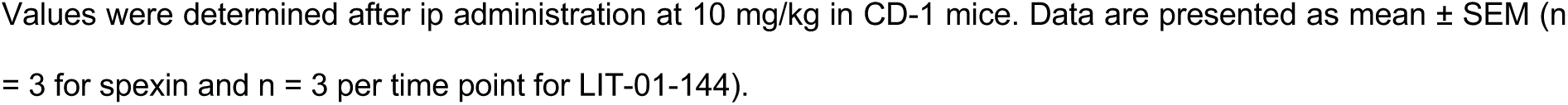
Pharmacokinetic parameters of spexin and LIT-01-144.

| Peptide | $C_{\max}$ (nmol/ml) | $T_{\max}$ (min) | $t_{1/2}$ (min) | AUC<sub>IP</sub> (min.nmol/ml) |
| --- | --- | --- | --- | --- |
| spexin | $0.13 \pm 0.05$ | 3 | $5.1 \pm 1.7$ | $0.74 \pm 0.06$ |
| LIT-01-144 | $0.20 \pm 0.02$ | 30 | $101 \pm 7$ | $35.3 \pm 3.8$ |

Values were determined after ip administration at 10 mg/kg in CD-1 mice. Data are presented as mean ± SEM (n = 3 for spexin and n = 3 per time point for LIT-01-144).

### Evaluation of the antinociceptive activity of spexin and LIT-01-144 after icv and systemic administration in naïve mice

We then evaluated the impact of icv administration of LIT-01-144 on the thermal nociceptive threshold of mice in comparison to spexin. As expected from previous experiments (Toll et al., 2012), spexin dose-dependently increased the nociceptive threshold of the animals (**Fig. 3A and 3B**). In the same experiment, administration of 0.1 and 1 nmol of LIT-01-144 elicited a significant antinociceptive response (**Fig. 3C and 3D**), demonstrating that LIT-01-144 possesses in vivo efficacy comparable to that of spexin, albeit at doses 10- to 100-fold lower, in accordance with our in vitro observations.

**Figure 3.**
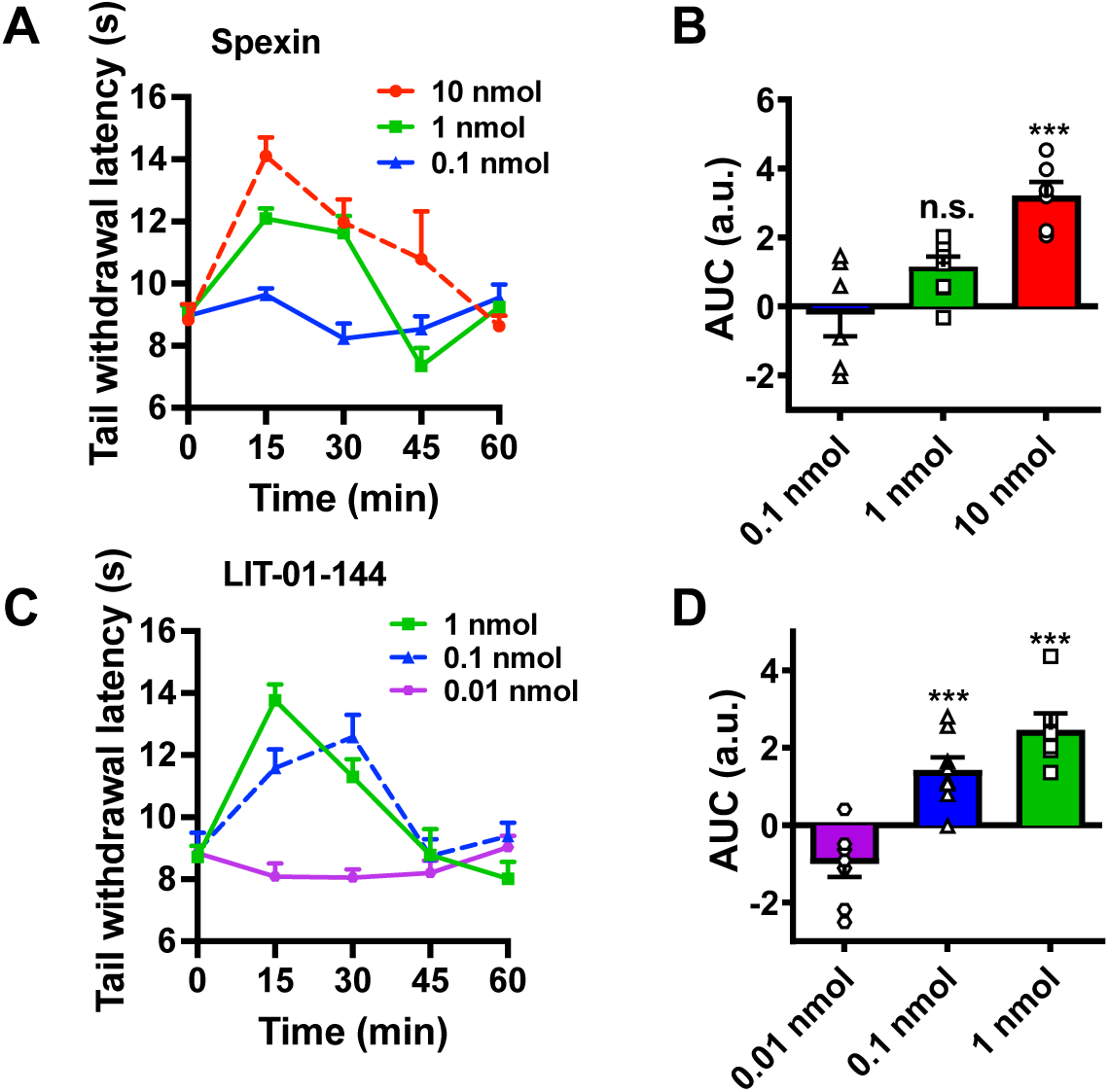
Antinociceptive effect of spexin and LIT-01-144 after central administration to naive mice. A, C) Time- and dose-dependent antinociceptive effect of spexin (A) and LIT-01-144 (C) in the tail immersion test (TIT) after icv administration in C57BL/6N mice. Withdrawal latencies are expressed in seconds and presented as mean ± SEM (n = 6-8 per group). B, D) Antinociceptive effect of spexin (B) and LIT-01-144 (D) in the TIT shown as AUC values over 60 min from A and C. Data are shown as individual values with means ± SEM (n = 6-8 per group). Groups were compared using one-way ANOVA followed by Bonferroni post hoc test ***p<0.001 compared to the dose of 0.1 nmol for spexin (B) and 0.01 nmol for LIT-01-144 (D).

We further studied the acute antinociceptive activity of LIT-01-144 and spexin following intraperitoneal (ip) administration in naïve mice at doses of 2 mg/kg and 20 mg/kg. We observed no significant effect of both compounds on thermal (**Fig. 4A**) and mechanical (**Fig. 4B**) nociceptive thresholds of the animals, suggesting that in pain free-mice, activation of peripheral GALR2/R3 do not induce antinociception.

**Figure 4.**
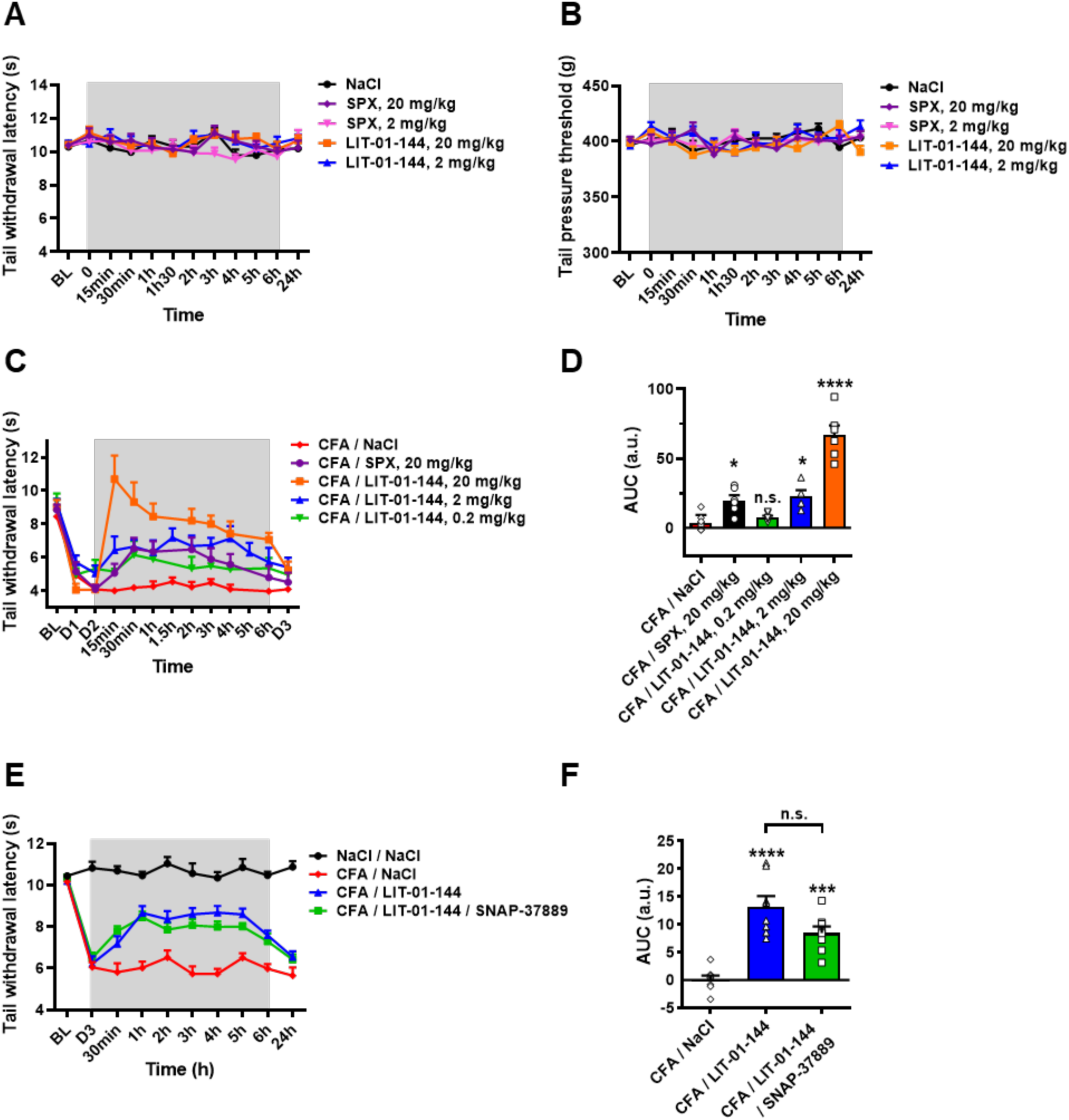
Antinociceptive effect of spexin and LIT-01-144 after systemic administration in mice. A, B) Effect of spexin and LIT-01-144 after ip administration at 2 and 20 mg/kg on the hot thermal (A) and mechanical (B) nociceptive thresholds of naive C57BL/6N mice in the TIT and the tail pressure test (TPT), respectively. Withdrawal latencies are expressed in seconds (A) and grams (B). Data are shown as means ± SEM (n = 7 per group). C) Time- and dose-dependent antinociceptive effect of spexin and LIT-01-144 in the TIT after ip administration in CFA-treated C57BL/6N mice. Withdrawal latencies are expressed in seconds and presented as means ± SEM (n = 6-8 per group). D) Total antinociceptive effect of spexin and LIT-01-144 in the TIT are shown as AUC values calculated from C, over 360 min. Data are shown as individual values with means ± SEM (n = 6-8 per group). E) Antinociceptive effect of LIT-01-144 (2 mg/kg, ip) in absence or presence of SNAP-37889 (0.33 mg/kg, ip) in CFA-treated C57BL/6N mice. Withdrawal latencies are expressed in seconds and shown as means ± SEM (n = 6-8 per group). F) Total antinociceptive effect LIT-01-144 in absence or presence of SNAP-37889 in the TIT are shown as AUC values calculated from E, over 360 min. Data are shown as individual values with means ± SEM (n = 6-8 per group). Groups were compared using one-way ANOVA followed by Bonferroni post hoc test *p<0.05, ***p<0.001, ****p<0.0001 compared to CFA/NaCl.

### Peripheral antinociceptive activity of spexin and LIT-01-144 in mice with inflammatory pain

We next evaluated the antinociceptive activity of spexin and LIT-01-144 after ip administration in mice with persistent inflammatory pain induced by sc administration of CFA in the tail. Conversely to what we observed in pain-free animals, both spexin and LIT-01-144 induced antinociception (**Fig. 4C and 4D**) when they were administered 2 or 3 days after CFA (when hyperalgesia was maximal). Similar to what we observed following icv administration, LIT-01-144 activity was significant at a dose that was 10-fold lower than spexin (**Fig. 4C and 4D**).

We then investigated which galanin receptor subtype was implicated in the antinociceptive response induced by LIT-01-144 in inflamed mice. To this end, we measured the activity of 2 mg/kg (ip) LIT-01-144 in the presence or absence of 0.33 mg/kg (ip) GALR3 selective antagonist SNAP-37889 (Swanson et al., 2005) on thermal nociception of CFA-mice. As shown in **Figures 4E and 4F** LIT-01-144 antinociceptive activity was similar to our previous observation (**Fig. 4C and 4D**) and was not affected by the GALR3 antagonist SNAP-37889 (**Fig. 4F**), indicating that the peripheral antinociceptive response of LIT-01-144 is mediated by GALR2 and not GALR3.

### Distribution of Galr2 transcript in DRG of naive and inflamed mice

In order to investigate the reason why LIT-01-144 displayed peripheral antinociceptive activity in inflamed but not in naïve animals, we studied the expression of *Galr2* transcript in DRG of naive vs inflamed mice by fluorescent *in situ* hybridization (ISH). In DRG from naïve animals, we observed a very low number of cells expressing *Galr2* transcript (**Fig. 5A**) while in DRG of CFA-mice we observed a higher number of *Galr2*-positive cells (**Fig. 5B and 5C**). Co-labeling of NeuN transcript showed that *Galr2* mRNA is mainly expressed in DRG neurons.

**Figure 5.**
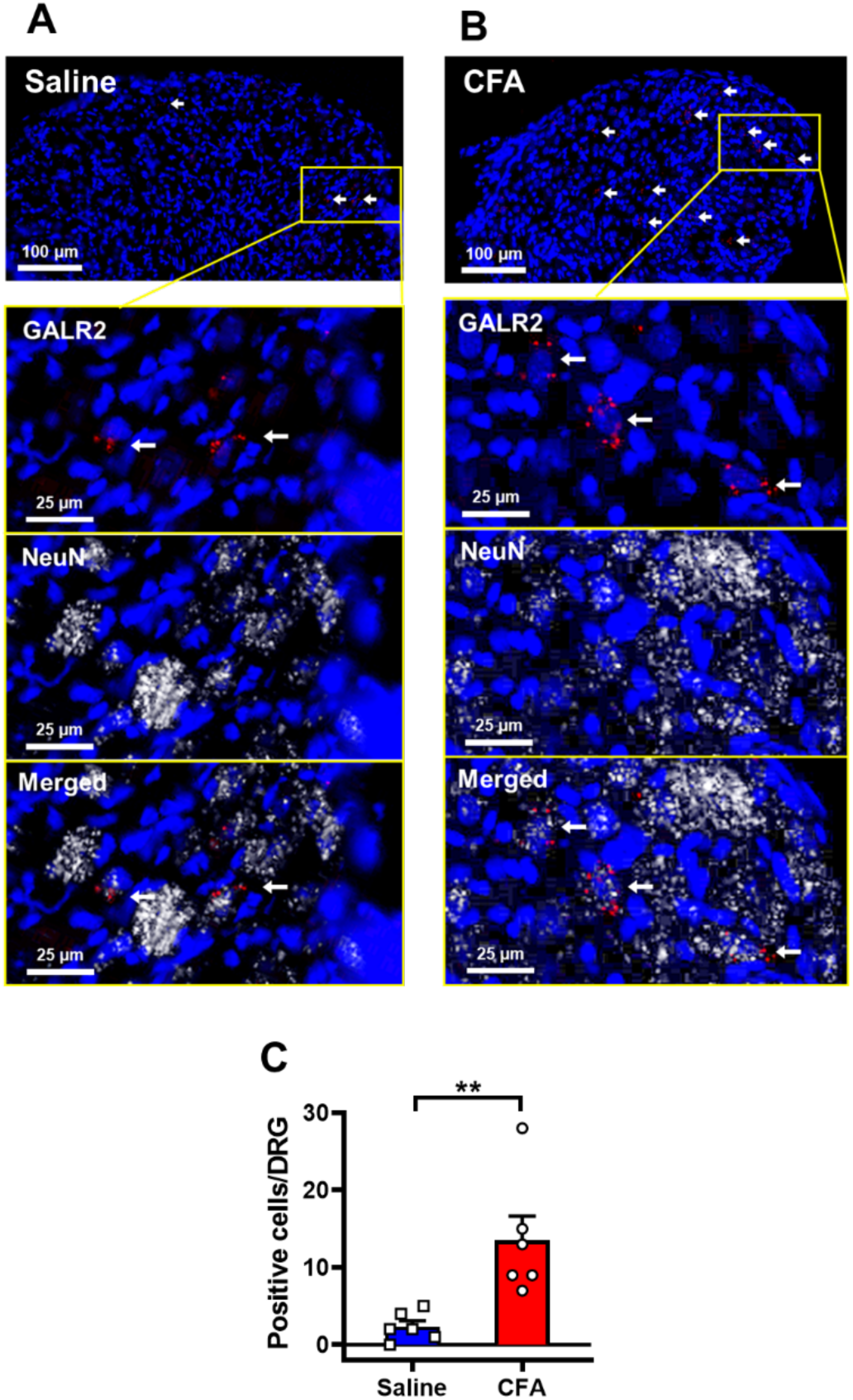
Distribution of *Galr2* mRNA in dorsal root ganglia (DRG) of normal and inflamed mice. A, B) Representative examples of cryosections of DRG from naïve (A) and CFA-treated mice (B) hybridized with a Cy3 fluorescent probe targeting *Galr2* transcript (red) and Cy5 fluorescent probe targeting NeuN transcript (white). Cells nuclei were stained with DAPI and are shown in blue. C) Comparison of the number of positive cells in DRG sections of inflamed mice vs normal mice (n = 6 per group). Groups were compared using unpaired parametric t-test **p<0.01.

We further investigated the distribution of *Galr2* transcript in mouse DRG neurons by using the publicly available single-cell RNA sequencing (scRNA-seq) atlas recently published (Bhuiyan et al., 2024). In naïve mouse DRG, *Galr2* is expressed predominantly in nociceptors (**Fig.6**). More specifically, Galr2 is found in a small part of both peptidergic (Calca-positive) and non-peptidergic (Mrgprd-positive) C-fiber neurons, which are responsible for sensing noxious stimuli that elicit pain reactions (**Fig.6B and 6C**). Altogether, these data suggest that LIT-01-144 antinociceptive peripheral activity in CFA-mice rely on inflammation-induced GALR2 expression in DRG nociceptors.

**Figure 6.**
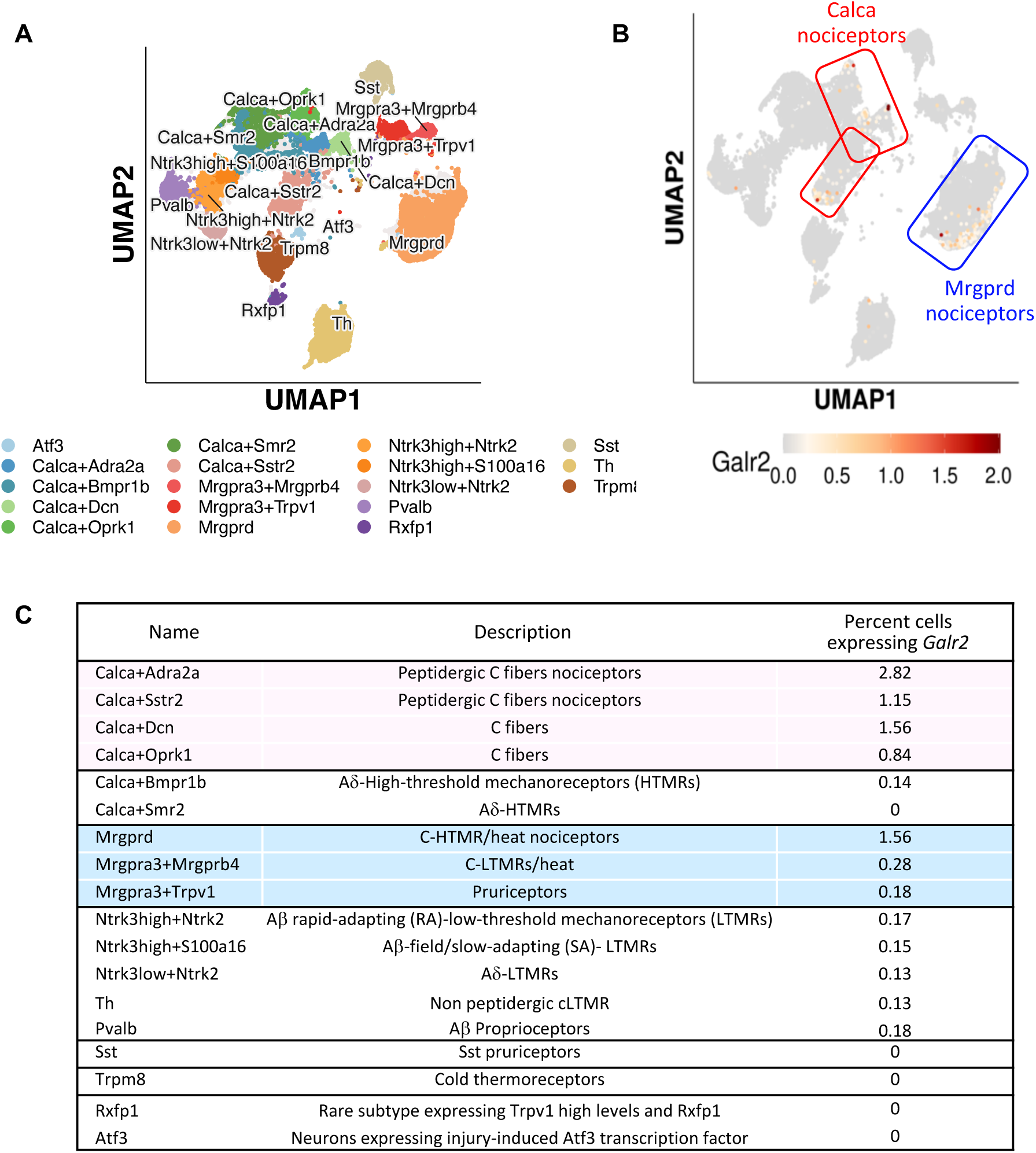
Distribution of *Galr2* mRNA in dorsal root ganglia (DRG) neuronal populations of normal mice. A) The recently published Harmonized cross-species atlases of trigeminal and dorsal root ganglia (DRG) has integrated single-cell RNA sequencing data-sets from five different mammalian species and enables to investigate the distribution of gene transcripts of interest in 18 distinct neuronal types (Bhuiyan et al., 2024). B) *Galr2* was found expressed in peptidergic (Calca-positive) and non-peptidergic (Mrgprd-positive) C-fiber nociceptors in the mouse datasets. C) The percents of each neuronal cell type expressing *Galr2* transcript are shown.

### Opioid dependency of antinociceptive activity of LIT-01-144

We further decided to evaluate the implication of the endogenous opioid system in the antinociceptive activity of LIT-01-144. CFA-mice were administered with LIT-01-144 at 2 mg/kg (ip) in absence or presence 1 mg/kg (sc) of naltrexone (**Fig. 7A**). As expected from our previous experiments (**Fig. 4C**), LIT-01-144 induced a significant antinociception as compared to saline treated animals, which was comparable to that observed in mice co-administered with naltrexone (**Fig. 7A and 7B**). These results are in agreement with previous results (Toll et al., 2012) and indicate that GALR2 mediated analgesic response is independent from the opioid system.

**Figure 7.**
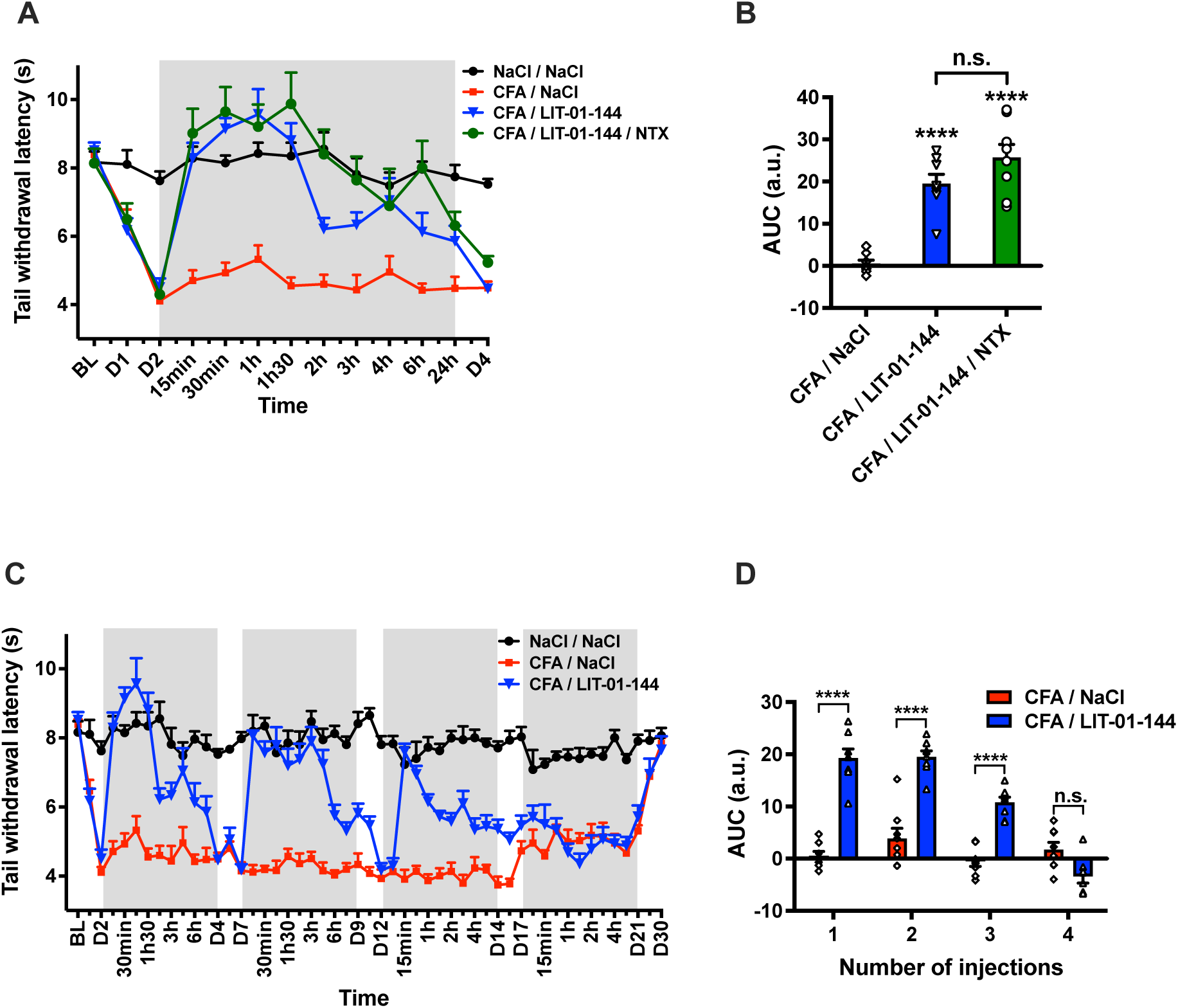
Opioid dependency of antinociceptive activity of LIT-01-144 and development of analgesic tolerance following repeated administrations in inflamed mice. A) Time-dependent antinociceptive effect of LIT-01-144 (2 mg/kg, ip) in absence or presence of naltrexone (1 mg/kg, sc) in the TIT in CFA treated C57BL/6N mice. Withdrawal latencies are expressed in seconds and shown as means ± SEM (n = 7-8 per group). B) Total antinociceptive effect of LIT-01-144 and LIT-01-144/naltrexone in the TIT shown as AUC values calculated from A, over 360 min. Data are shown as individual values with means ± SEM (n = 7-8 per group). Groups were compared using one-way ANOVA followed by Bonferroni post hoc test ****p<0.0001 compared to CFA/NaCl. C) Development of analgesic tolerance upon chronic treatment with LIT-01-144. C57BL/6N mice received injections of LIT-01-144 (2 mg/kg, ip) or saline on days 2, 7, 12 and 17. Data are expressed as means ± SEM (n= 7-8 per group). D) Total antinociceptive effect of LIT-01-144 and LIT-01-144/naltrexone in the TIT shown as AUC values calculated from C, over 360 min. Data are shown as individual values with means ± SEM (n= 7-8 per group). For each injection, groups were compared with the CFA-saline control group using unpaired parametric t-test ****p<0.0001.

### Development of analgesic tolerance following LIT-01-144 repeated administrations

In order to study whether LIT-01-144 could induce analgesic tolerance, we performed repeated administrations of LIT-01-144 (2 mg/kg, ip) in CFA-mice. As shown in **Figure 7C and 7D**, the first administration of LIT-01-144 on day 2 post-CFA induced a significant increase of tail withdrawal latency of the animals that lasted at least 6 hours and returned to values observed in mice injected with CFA alone on day 4. The second administration on day 7 induced a similar increase of tail withdrawal latencies that returned to CFA control values only on day 12. The amplitude of the antinociceptive activity following the third LIT-01-144 administration on day 12 appeared to be lower as compared to previous administrations, and no significant antinociception was observed after the last LIT-01-144 administration on day 17 (**Fig. 7C and 7D)**. Both CFA- and CFA + LIT-01-144-treated animals returned to basal nociceptive values on day 30. These data indicate that, similarly to what is observed with opioids, repeated activation of GALR2 with LIT-01-144 induces analgesic tolerance.

## Discussion

In this study, we report the successful design and *in vitro* and *in vivo* characterization of a metabolically stable GALR2/3 agonist LIT-01-144. *In vitro*, LIT-01-144 displays potent agonist activity at GALR2/R3 while retaining the same selectivity than spexin toward GALR1. *In vivo*, LIT-01-144 displays increased metabolic stability as compared to spexin and antinociceptive activity at low doses when administered at the central level (icv) and after peripheral administration (ip) in mice with persistent inflammatory pain. This antinociceptive activity was specifically mediated through GALR2 and was independent of opioid receptor pathways. Taken together, these results establish LIT-01-144 as a valuable pharmacological tool for investigating GALR2-driven mechanisms involved in the modulation of pain and other functions and underscore GALR2 as a compelling peripheral target for the development of non-opioid analgesics.

Developing pharmacological tools derived from endogenous peptides is fundamental for elucidating the involvement of a biological target in a given pathology. The main limitation to their use *in vivo* is their lack of metabolic stability and numerous strategies have been developed over the last decades to address this issue (Ma et al., 2025). Among them, amino acid substitution is a well-known approach for improving the pharmacokinetic (PK) properties of peptide-based pharmacological tools. However, this approach entails the replacement of residues, for instance with D-amino acids, within the core sequence, which can compromise both biological activity and selectivity (Fosgerau and Hoffmann, 2015). Similarly, peptide cyclization, whether through side-chain linkages within the sequence or via N- to C-terminal connections, can significantly influence the conformation, potentially leading to diminished activity and selectivity (Muttenthaler et al., 2021). In contrast, peptide concealment offers a promising alternative. Exopeptidic moieties (lipid, PEG …) can be readily introduced at the N-terminus or on side chains with a lower risk of disrupting the peptide biological activity or pharmacological profile. Nonetheless, current peptide concealment techniques present certain limitations. For example, PEG chains have been implicated in potential nephrotoxicity and immunogenic responses (Zhang et al., 2016; Hoang Thi et al., 2020; Kozma, 2020). We describe here a novel alternative to develop a metabolically stable derivative of spexin. This approach involves the site-specific incorporation of a fluorocarbon moiety into spexin amino acid sequence. This methodology has previously enabled the development of highly potent, selective and metabolically stable apelin peptides for the *in vivo* study of apelin receptor as a potential target for the treatment of hyponatremia, multiple sclerosis or cardiovascular disorders (Gerbier et al., 2017; Flahault et al., 2021; Birmpili et al., 2024; Girault-Sotias et al., 2024). In this study, we applied the same strategy to spexin by incorporating a FC at both the N-terminal and C-terminal positions, yielding two fluorinated spexin derivatives: LIT-01-144 and LIT-01-128, respectively. The N-terminal position was shown to be the most suitable to modify spexin, resulting in a potent and stable probe *in vitro*. These findings are consistent with previously reported N-terminal modifications of spexin (Fmoc, PEG, pyroglutamyl), demonstrating the permissiveness of its N-terminus to chemical modification (Reyes-Alcaraz et al., 2016). Of particular importance, the incorporation of FC not only did not hamper the interaction of the peptide with GALR2 and GALR3 without modifying the selectivity toward GALR1 but also enabled a 10-fold increase of the functional activity of LIT-01-144 at GALR2 and GALR3 as compared to spexin. Additionally, as expected, the presence of FC on LIT-01-144 resulted in a significant improvement of *in vitro* and *in vivo* stability leading to a strong increase of mice exposition as compared to spexin at the same dose. Consistently with *in vitro* results, LIT-01-144 antinociceptive activity was observed at doses that were 10- to 100-times lower than spexin in naïve mice after central administration and in inflamed mice after systemic administration. This achievement highlights the versatility of the FC modification method, which we employed to modify an endogenous peptide targeting a new family of receptors, GALRs, and addressing a novel pathological context–pain. These results are consistent with previous reports on peptide modifications by FC (Gerbier et al., 2017; Flahault et al., 2021; Birmpili et al., 2024; Girault-Sotias et al., 2024). Moreover, they represent the first example of a successful FC modification that enhances both *in vitro* and *in vivo* activity of a neuropeptide. Altogether, our data point to LIT-01-144 as a suitable tool to study the role of GALR2/R3 *in vivo* and further support the broader applicability of this modification approach to other endogenous peptides whose limited metabolic stability has impeded the investigation of their physiological roles.

Pain has long been a central focus of biomedical research, with considerable efforts devoted to developing alternatives to opioids for pain management. Nevertheless, opioids remain the gold standard for analgesia, underscoring the urgent need for next-generation pain therapeutics (Lyden and Binswanger, 2019). The well-characterized galanin receptor (GALR) family has been implicated in pain mechanisms across various species (Lang et al., 2015; Fonseca-Rodrigues et al., 2022). However, the lack of selective and metabolically stable ligands has limited both the investigation of GALRs specific roles in pain modulation and the development of therapeutic candidates targeting these receptors (Webling et al., 2012). This is particularly true for GALR2 for which contradictory studies reported either a pronociceptive (Jimenez-Andrade et al., 2004, 2006; Chen et al., 2018) or antinociceptive (Metcalf et al., 2015; Zhang et al., 2017) activity. These contrasting results were suggested to be due to differences in the pain models used in these studies but the lack of selectivity of the ligands used to study the galanin receptors can also be questioned (Webling et al., 2012). Moreover, the absence of robust GALR-KO mouse models, particularly GALR2 KO mice that display impaired development of DRG neurons (Hobson et al., 2006) does not allow to confirm the results obtained using the poorly selective galanin ligands. In this context, LIT-01-144 emerges as a highly selective ligand for GALR2 and GALR3, with almost no activity at GALR1. Moreover, its peripheral antinociceptive activity was not blocked by the GALR3 selective antagonist SNAP-37889 (Swanson et al., 2005), nor by the opioid antagonist naltrexone. These observations provide novel evidence that GALR2 and its endogenous ligand (spexin and/or galanin) constitute a *bona fide* antinociceptive system independent of the opioid system.

In this study we also show that the number of *Galr2*-positive neurons is low in DRG of naïve mice but is strongly increased in DRG of inflamed mice. This observation is in agreement with our behavioral data showing no activity of peripheral administration of LIT-01-144 at high dose in naïve animals while it displayed potent antinociceptive action in inflamed animals. These data are in good accordance with previous report showing increase of labelling intensity and number of Galr2 mRNA positive neurons in rats with inflammatory pain (Sten Shi et al., 1997), increase of GALR2 immunoreactive cells in rats with neuropathic pain (Chen et al., 2018) as well as increase in number and intensity of GALR2-EGFP positive cells in mice with neuropathic pain (Lyu et al., 2020). In addition, mining of single cell RNA-seq databases (Bhuiyan et al., 2024) confirmed the expression of *Galr2* in a small number of DRG neurons from naïve mice that are specifically involved in the modulation of nociception including peptidergic and non-peptidergic neurons. A similar observation was made in human DRG (Yu et al., 2024). Altogether these data suggest that GALR2 could represent an interesting target for chronic pain treatment.

In conclusion, our study together with our previous observations with apelin (Gerbier et al., 2017; Flahault et al., 2021; Birmpili et al., 2024; Girault-Sotias et al., 2024) outline the potential of adding FC on peptides to develop pharmacological tools from endogenous peptide sequence and allowed us to provide novel evidence that GALR2 displays peripheral antinociceptive properties in persistent pain and represent an interesting target to develop novel analgesics for the treatment of chronic inflammatory pain.

## Acknowledgments

This work was supported by the the Agence Nationale de la Recherche (ANR-16-CE18-0030), CNRS, University of Strasbourg, SATT Conectus, LABEX Medalis (ANR-10-LABX-0034, Programme d’investissement d’avenir), Graduate School of Pain EURIDOL of the University of Strasbourg (ANR-17-EURE-0022, Programme d’investissement d’avenir), and by the SFRISTRAT’US Project (ANR-20-SFRI 0012) and IdEx Unistra (ANR-10-IDEX-0002) under the framework of the French Program “Investments for the Future” given to the Strasbourg Drug Institute (IMS), as part of the Interdisciplinary Thematic Institute (ITI) 2021−2028 program of the University of Strasbourg, CNRS and Inserm. The authors thank the PACSI platform for technical assistance.

## Abbreviations

ANOVA: Analysis Of Variance
AUC: Area Under the Curve
BSA: Bovine Serum Albumin
cAMP: Cyclic Adenosine Monophosphate
CFA: Complete Freund’s Adjuvant
CNS: Central Nervous System
DMEM: Dulbecco’s Modified Eagle Medium
DRG: Dorsal Root Ganglia
FC: Fluorocarbon Chain
GALR: Galanin Receptor
GPCR: G protein-coupled receptor
HEK: Human Embryonic Kidney
hGal: Human Galanin
icv: Intracerebroventricular
ip: Intraperitoneal
ISH: In situ hybridization
KO: Knockout
LC-MS: Liquid Chromatography – Mass Spectrometry
RP-HPLC: Reversed-Phase High-Performance Liquid Chromatography
sc: Subcutaneous
t_1/2_: Half-life time
TIT: Tail Immersion Test
TPT: Tail Pressure Test
TSA: Tyramide Signal Amplification
UPLC-MS/MS: Ultra Performance Liquid Chromatography tandem Mass Spectrometry

## Notes

### Competing Interest Statement

The authors have declared no competing interest.

