## Supplemental Table 1 and Sup methods for "A metabolically resistant spexin analogue, LIT-01-144, induces potent non-opioid peripheral antinociception in persistent pain via activation of GALR2"

#### Supplementary Material

**Table S1** Amount of spexin and LIT-01-144 in CD1-mouse brain following IP administration at 10 mg/kg after 15, 30 and 360 min.

| Time (min) | Brain concentration (nmol/g of brain) | |
|---|---|---|
| | Spexin | LIT-01-144 |
| 5 | < 0.008 | < 0.1 |
| 30 | < 0.008 | < 0.1 |
| 360 | < 0.008 | < 0.1 |

Brain concentration of spexin and LIT-01-144 were determined by LC-MS/MS using a calibration curve. 3 different mice were used for each timepoint and molecule.

##### Supplementary methods for chemistry

##### Synthesis

##### General information

Reagents were obtained from commercial sources and used without any further purification. Fmoc-L-amino acids were purchased from Iris Biotech. Fmoc-protected preloaded Rink-AM resin was purchased from Sigma-Aldrich and the overall yields for the solid-phase syntheses were calculated based on the initial loadings provided by the supplier (0.4 $mmol \cdot g^{-1}$). Reactions were monitored by analytical reverse-phase high-performance liquid chromatography (RP-HPLC), performed on a C18 Kinetex column (5 µm, 4.6 mm × 150 mm) using a linear gradient (5% to 95% in 7.39 min, flow rate of 1.5 $ml \cdot min^{-1}$) of MeCN with 0.1% TFA (v/v) in $H_2O$ with 0.1% TFA (v/v) with detection at 220 and 254 nm. High-resolution mass spectra (HRMS) were obtained on an Agilent Technologie 6520 Accurare-Mass Q.Tof LC/MS apparatus equipped with a Zorbax SB C18 column (1.8 µm, 2.1 × 50 mm) using electrospray ionization (ESI) and a time-of-flight analyzer (TOF).

##### Automated solid phase peptide synthesis (SPPS)

Automated SPPS was carried out using a Liberty Blue synthesizer (CEM) by standard Fmoc solid-phase peptide chemistry on a preloaded Rink-AM resin (0.4 $mmol.g^{-1}$ resin, 0.1 mmol scale), using DMF as solvent. The coupling of each amino acid (2 M, 10 eq.) was carried out using DIC (0.5 M) and Oxyma (1 M). Fmoc groups were removed using a 20% (v/v) solution of piperidine in DMF. Washing steps were performed using DMF. At the end of peptide synthesis, the resin was washed with DCM, MeOH, and $Et_2O$, then dried under a high vacuum.

**Manual solid phase peptide synthesis (SPPS)**

Manual SPPS was performed in polypropylene tubes equipped with polyethylene frits and polypropylene caps using an orbital agitator shaking device starting from the dried resins obtained by automated SPPS. The Fmoc-protected resin (1 eq.) was swollen for 1 h in DCM and the excess of solvent was removed by filtration. The N-terminal Fmoc group was removed by using a 20% (v/v) solution of piperidine in DMF (2 times for 15 min). All Fmoc cleavage steps were performed in the same way. The piperidine solution was drained off and the resin was washed three times with successively DMF, DCM, and MeOH. All Fmoc-protected amino acids (4 eq.) were coupled in DMF (7 ml per 0.1 mmol of resin) for 45 min using HATU (3.9 eq.) in the presence of DIPEA (12 eq.) as activating agent. 4,4,5,5,6,6,7,7,8,8,9,9,10,10,11,11,11-Heptadecafluoroundecanoic acid (4 eq.) was coupled in DMF (7 ml per 0.1 mmol of resin) for 45 min using HATU (3.9 eq.) in the presence of DIPEA (12 eq.) as activating agent. The excess of solvent was removed by filtration and the resin was washed three times with successively DMF, DCM, and MeOH.

**General procedure for peptide acetylation**

Deprotected peptides were acetylated using a solution of acetic anhydride (10%), DIEA (5%) in DCM. The solution (1 ml/100 mg of resin) was added and the syringe was stirred using an orbital agitator for 10 min. The acetic anhydride solution was drained off under vacuum and the resine was washed three times using DCM.

**Resin cleavage and peptide purification**

Peptides were cleaved from the resin under classical cleavage conditions with TFA/phenol/$H_2O$/thioanisol/EDT 80/7.5/5/5/2.5 (v/v) (7 ml per 0.1 mmol of resin). The mixture was stirred at room temperature for 3 h. The solution was filtered under vacuum and the peptides were precipitated with cold diethyl ether (30 volumes per volume of the cleavage mixture). Precipitated peptides were centrifuged at 4°C at 3000 g for 2 min. The diethyl ether solution was removed by decantation and the crude solid peptides were dried *in vacuo*. Crude peptides were purified by semi-preparative reversed-phase HPLC chromatography (Gilson PLC2020 system) on a SunFire C18 column (5 µm, 19 × 150 mm) using a linear gradient (5% to 95% in 40 min, the flow rate of 18 ml.min$^{-1}$) of MeCN with 0.1% TFA (v/v) in $H_2O$ with 0.1% TFA (v/v). Detection was set at 220 and 254 nm. Fractions containing the desired peptide were collected and freeze-dried.

**General procedure for peptide analysis**

RP-HPLC was performed on an Agilent Technologies 1260 Infinity II HPLC system with a Kinetex EVO C18 column (2.7 µm, 4.6 mm × 150 mm) using a linear gradient (5% to 100% in 20 min, flow rate of 1.5 ml/min ) of solvent B (0.1% TFA in ACN, v/v) in solvent A (0.1% TFA in H2O, v/v). Detection was set at 220.4 and 254.4 nm.

**Synthetic procedures**

**LIT-01-128**

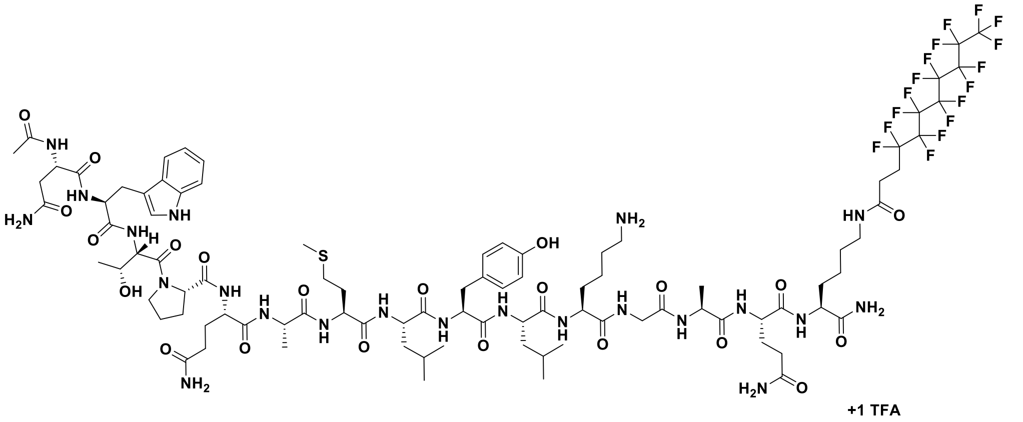

Fmoc-N(Trt)W(Boc)T(tBu)PQ(trt)AMLY(tBu)LK(Boc)GAQ(Trt)K(Mmt)-Rink sequence was synthesized following the general method for automated SPPS (0.1 mmol scale). K15(Mmt) was selectively deprotected using a solution of DCM/TIS/TFA (94/5/1 v/v) (7 times for 2 min). The 1% TFA solution was drained off and the resin was washed three times using DCM. 4,4,5,5,6,6,7,7,8,8,9,9,10,10,11,11,11-Heptadecafluoroundecanoic acid was introduced on the resin-bound peptide (28 µmol) following the general method for manual SPPS. The peptide was N-terminally deprotected and capped following general procedure for peptide acetylation. The peptide was cleaved and purified following the general method, affording the title compound as white solid (19.4 mg, 29%). Analytical HPLC $t_R$ = 9.91 min (> 95% purity at 220 nm and 254 nm); **HRMS** (ESI-TOF): Calculated for $C_{93}H_{133}F_{17}N_{22}O_{22}S$ $[M+2H]^{2+}/2$: 1132.4707; found: 1132.4726 ($\Delta_{M/Z}$ = 1.67 ppm).

**LIT-01-144**

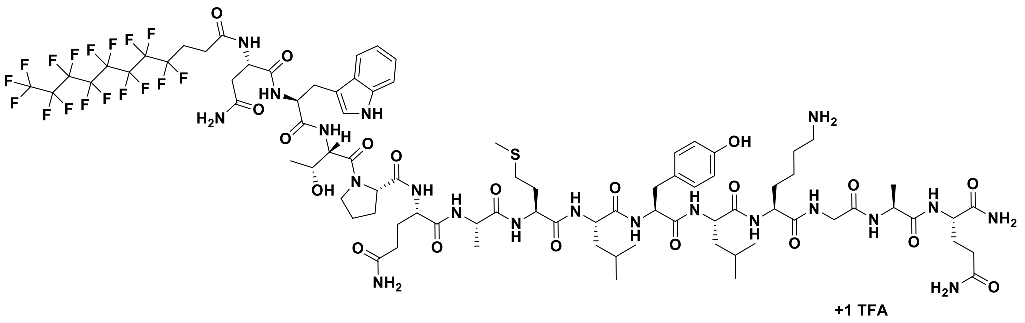

Fmoc-N(Trt)W(Boc)T(tBu)PQ(trt)AMLY(tBu)LK(Boc)GAQ(Trt)-Rink sequence was synthesized following the general method for automated SPPS (0.1 mmol scale). 4,4,5,5,6,6,7,7,8,8,9,9,10,10,11,11,11-Heptadecafluoroundecanoic acid was introduced on the resin-bound peptide (72 µmol) following the general method for manual SPPS. The peptide was cleaved and purified following the general method, affording the title compound as white solid (11.9 mg, 8%). Analytical HPLC $t_R$= 10.08 min (>95% purity at 220 nm and 254 nm); **HRMS** (ESI-TOF): Calculated for $C_{85}H_{119}F_{17}N_{20}O_{20}S$ $[M+2H]^{2+}/2$: 1047.4179; found: 1047.4192 ($\Delta_{M/Z}$ = 1.24 ppm).

### HRMS analyses

#### LIT-01-128

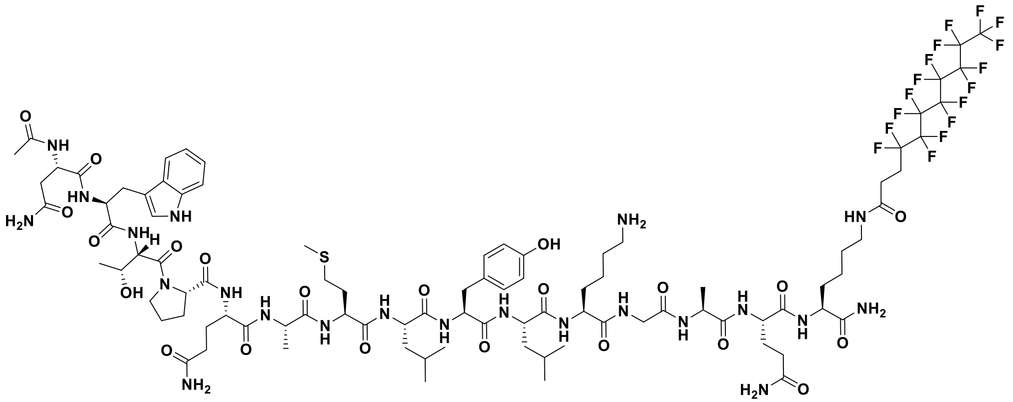

+1 TFA

Fragmentor Voltage 180 Collision Energy 0 Ionization Mode ESI

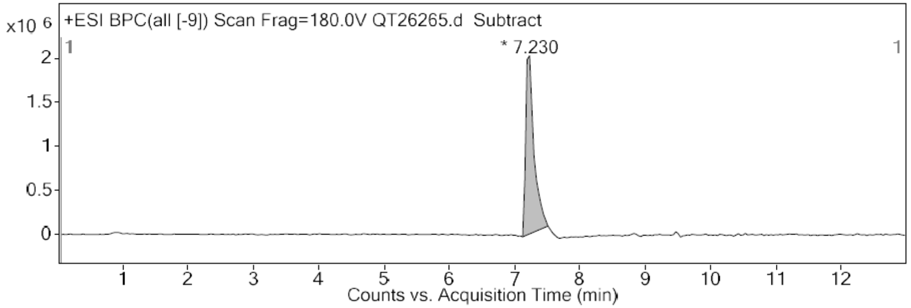


**Integration Peak List**

| Peak | Start | RT | End | Height | Area | Area % | AreaSum% |
|---|---|---|---|---|---|---|---|
| 1 | 7.123 | 7.23 | 7.514 | 2017833.29 | 18079349.73 | 100 | 100 |

Fragmentor Voltage 180 Collision Energy 0 Ionization Mode ESI

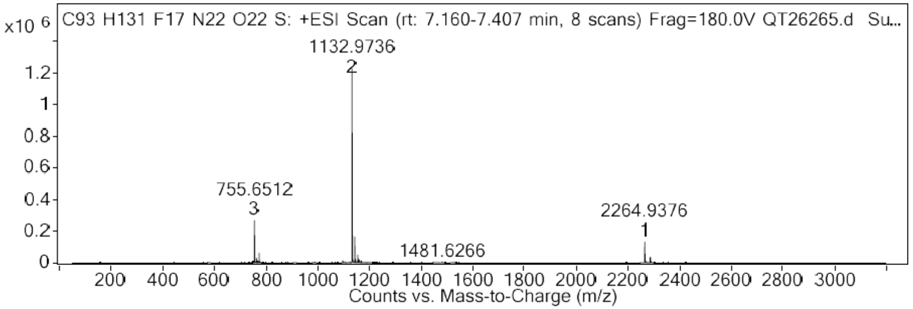


**Peak List**

| *m/z* | z | Abund | Formula | Ion |
|---|---|---|---|---|
| 755.3172 | 3 | 229646.5 | | |
| 755.6512 | 3 | 268397.4 | | |
| 755.9848 | 3 | 174448.1 | | |
| 1132.4726 | 2 | 1072310.52 | C93 H131 F17 N22 O22 S | (M+2H)+2 |
| 1132.9736 | 2 | 1236918.04 | C93 H131 F17 N22 O22 S | (M+2H)+2 |
| 1133.474 | 2 | 827707.11 | C93 H131 F17 N22 O22 S | (M+2H)+2 |
| 1133.975 | 2 | 416574.83 | C93 H131 F17 N22 O22 S | (M+2H)+2 |
| 1134.4752 | 2 | 170232.94 | C93 H131 F17 N22 O22 S | (M+2H)+2 |
| 1143.4614 | 2 | 139407.18 | | |
| 1143.9627 | 2 | 161702.94 | | |

**LIT-01-144**

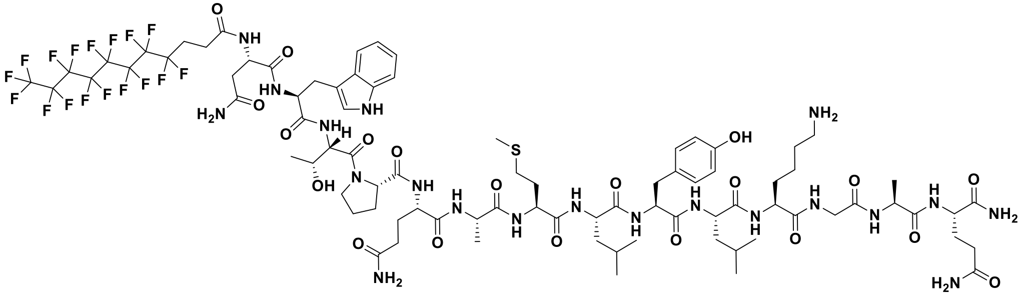

+1 TFA

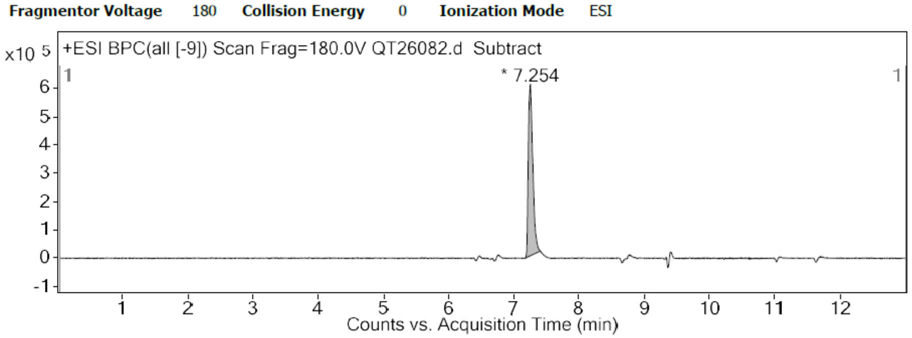


**Integration Peak List**

| Peak | Start | RT | End | Height | Area | Area % | AreaSum% |
|---|---|---|---|---|---|---|---|
| 1 | 7.179 | 7.254 | 7.406 | 606319.77 | 2943021.55 | 100 | 100 |

**Peak List**

| *m/z* | z | Abund | Formula | Ion |
|---|---|---|---|---|
| 698.6149 | 3 | 74697.63 | | |
| 698.9487 | 3 | 79652.55 | | |
| 699.2822 | 3 | 48115.29 | | |
| 1047.4192 | 2 | 354386.86 | | |
| 1047.9204 | 2 | 376577.84 | | |
| 1048.4211 | 2 | 235002.54 | | |
| 1048.9214 | 2 | 110538.43 | | |
| 1058.4087 | 2 | 67467.33 | | |
| 1058.91 | 2 | 71431.23 | | |
| 2094.8325 | 1 | 45254.41 | C85 H117 F17 N20 O20 S | (M+H)+ |

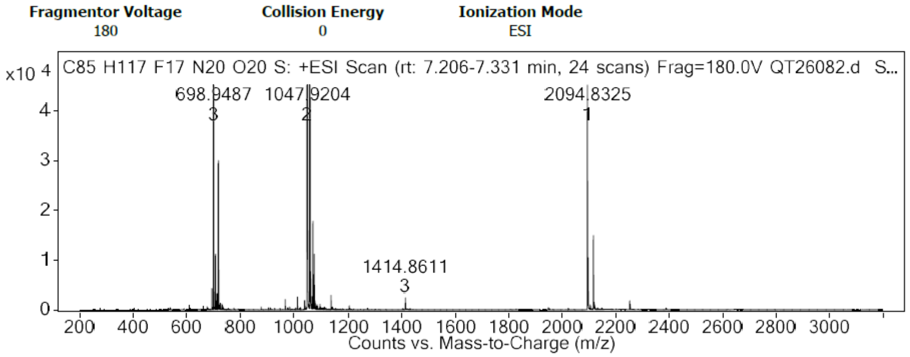
